# Biochemical and metabolic maladaption defines pathological niches in progressive multiple sclerosis

**DOI:** 10.1101/2022.09.26.509462

**Authors:** Melissa Grant-Peters, Charlotte Rich-Griffin, Hing-Yuen Yeung, Tom Thomas, Simon Davis, Mohammad Azizian, Joseph Fisher, Roman Fischer, Gianfelice Cinque, Calliope A. Dendrou

**Affiliations:** Nuffield Department of Medicine, Wellcome Centre for Human Genetics, University of Oxford, Oxford, UK; The Kennedy Institute of Rheumatology, University of Oxford, Oxford, UK; Translational Gastroenterology Unit, NIHR Oxford Biomedical Research Centre, Oxford University Hospitals NHS Foundation Trust, John Radcliffe Hospital, Oxford, United Kingdom; Target Discovery Institute, Nuffield Department of Medicine, University of Oxford, Oxford, UK; Diamond Light Source, Didcot, UK

## Abstract

Progressive multiple sclerosis (MS) is driven by demyelination, neuroaxonal loss, and mitochondrial damage occurring behind a closed blood-brain barrier (BBB).^1,2^ Patients with progressive MS typically fail to respond to available immunomodulatory drugs that reduce relapses in early disease.^2^ This indicates a dire need to identify non-canonical therapeutic avenues to limit neurodegeneration and promote protection and repair.^3^ Here, we have employed high-resolution multiomic profiling to characterise the biochemical and metabolic adaptations underpinning MS pathology, as these have been incompletely described but critically, may be amenable to BBB-permeable drug targeting. Using synchrotron radiation (SR)- and focal plane array (FPA)-based Fourier transform infrared microspectroscopy (μFTIR), we spatially mapped the biochemical features present in human progressive MS and control post-mortem brain and rare spinal cord tissue. By employing single-nuclear RNA sequencing (snRNA-seq), 10x Genomics Visium spatial transcriptomics and spatial proteomics to resolve their cellular context, we found that these biochemical features provide a uniquely and highly disease-specific barcode for distinct pathological niches within the tissue. Characterisation of the metabolic processes underpinning these niches revealed an associated re-organisation of the astrocytic landscape in the grey and white matter, with implications for the treatment of progressive MS.

---

FTIR microspectroscopy is emerging as a label-free and non-destructive technology for the high-resolution spatial mapping of biochemical signatures present within biological samples,^4,5^ but has never been applied to the analysis of CNS tissue from MS patients. We employed SR-μFTIR utilizing the Diamond Light Source MIRIAM beamline with a 0.964 cm^−1^ spectral and a 5×5 μm^2^ spatial resolution, to obtain 2,061 spectra from 10 μm-thick brain and spinal cord sections from our secondary progressive MS (SPMS) patient cohort (**Fig. 1**). By exciting the tissue with incident light within the infrared wavelength range, sample absorbance characteristics were captured as different vibrational modes corresponding to specific functional groups present in lipids, proteins, nucleic acids, and carbohydrates (**Fig. 2a**). The spectra were obtained from areas within each tissue section that were characterised as lesional and normal-appearing white matter (NAWM) through histological staining, and the variance in the data generated was explored through dimensionality reduction. We found that the highest proportion (60.12%) of the observed variance (principal component (PC) 1) was predominantly driven by the absorbances detected in the 2,800-3,000 cm^−1^ wavenumber region (**Fig. 2b**). Absorbance peaks within this region correspond to the presence of lipids (**Fig. 2a**), and spectra derived from the WM lesion areas showed a significantly lower infrared light absorbance than NAWM spectra (**Fig. 2c**), consistent with the demyelinating nature of MS pathology.^1^ PC2, which explained 8.37% of the variance in the data, was driven by absorbances detected in the 2,800-3,000 cm^−1^ and also in the 1,500-1,700 cm^−1^ regions (**Fig. 2b**); the latter contains peaks corresponding to the amide groups found in proteins (**Fig. 2a**). In contrast to the lipid-associated spectral peaks, the absorbance of the amide peaks was significantly higher in the WM lesion compared to the NAWM spectra (**Fig. 2c**). Notably, in addition to providing information on overall protein content, the shape of the amide peaks enables inference of protein secondary structure motifs, such that the relative proportion of *α*-helices, *β*-sheets and *β*-turns can be estimated.^6^ Alterations in the proportions of these protein secondary structure features can be indicative of proteinopathy: in Alzheimer’s disease *β*-sheet content is elevated in amyloid beta plaques^7^ and in Parkinson’s disease increased *β*-sheet content has been detected in neurons of the substantia nigra.^8^ Interestingly, we found significantly higher percentages of *β*-sheets and *β*-turns and a lower percentage of *α*-helices in the WM lesions relative to the NAWM (**Fig. 2c**). This provides direct evidence of altered protein folding in SPMS lesions, and suggests that deviation from protein homeostasis is a pathophysiological mechanism common to diseases with a neurodegenerative component, regardless of their etiology.

Although the PC analysis of the SR-μFTIR data demonstrated that the top PCs could distinguish subsets of spectra from the WM lesion and NAWM areas, it also revealed additional substructure in the data (**Fig. 2b**). Therefore, in order to better resolve the biochemical heterogeneity of the patient tissue, we employed FPA-μFTIR with a globar source to scan more continuous areas, spanning demyelinated and non-demyelinated WM as well as grey matter (GM) areas, and to profile CNS tissue from controls (**Fig. 1**). We obtained a total of 229,376 spectra with a 1.928 cm^−1^ spectral and a 5×5 μm^2^ spatial resolution. Implementing an unsupervised dimensionality reduction approach,^9,10^ we identified seven spectral clusters, five of which (termed Les1-Les5) were almost exclusively derived from SPMS tissue, collectively comprising <0.5% of the control spectra (**Fig. 2d**). The other two clusters corresponded to healthy WM and GM spectra and were detected in both the control and case tissue. Inspection of the spectral cluster features showed that each had a unique biochemical fingerprint characterized by variation in the lipid and protein content, protein secondary structure and the carbonyl ester content (1,700-1,760 cm^−1^ region), which can be used to derive a metric of lipid oxidation^11^ (**Fig. 2e, Supp. Fig. 1**). Oxidized lipids have been previously reported to be present in MS lesions,^12^ and we observed that SPMS-specific clusters with high lipid oxidation also had a high *β*-sheet and *β*-turn content, and this was most prominent in the GM (Les4). Notably, whilst under physiological conditions protein folding requires redox processes, under conditions of excessive oxidation proteins can become carbonylated resulting in alterations to their conformation and even aggregation.^13^ To ascertain how the biochemomic fingerprint of the different FPA-μFTIR clusters related to the underlying cellular architecture of the tissue, we performed mass spectrometry-based spatial proteomics on 89 samples of 316×316 μm^2^ resolution from consecutive/near-consecutive 10 μm tissue sections (**Fig. 1**). Spatially aligning the FPA-μFTIR clusters and proteomics data histologically, we first sought to confirm the relationship between FPA-μFTIR cluster lipid content and the expression of myelin basic protein (MBP), which is characteristically lost in MS in demyelination. As anticipated we found that MBP expression was highest in tissue samples that corresponded to the healthy WM spectral cluster, and in fact cluster lipid content was highly positively correlated with MBP level (*r*=0.982, *P*=2.573×10^−3^; **Supp. Fig. 2a**), despite the lipid content being inversely correlated with the overall protein content (*r*=-0.919, *P*=3.432×10^−3^; **Supp. Fig. 2b**). Next, we performed unsupervised clustering of the spatial proteomics data. This revealed that the FPA-μFTIR clusters largely segregated with the proteomics clusters (**Supp. Fig. 2c**). For example, SPMS-specific cluster Les1 predominantly co-segregated with proteomics cluster 2 which was enriched for proteins corresponding to oligodendrocytes (e.g. MBP, PLP1), and to a lesser extent astrocytes (e.g. GFAP, AQP4, HEPACAM, C3) and immune cells (e.g. IGHG1); cluster Les2 was more associated with astrocyte and immune proteins in proteomics cluster 4; and clusters Les4 and Les5 were verified as GM given their association with neuronal proteins (e.g. SYN1, CAMK2B) in proteomics cluster 2. To further confirm these findings spatially at the RNA level (**Fig. 2f**), we generated the first higher resolution spatial transcriptomic data of SPMS postmortem tissue by using the 10x Genomics Visium platform in conjunction with Cell2location-based deconvolution (**Fig. 1**). The deconvolution was performed by leveraging a brain snRNA-seq atlas of 71,078 nuclei derived from the integration of our multiplexed 3’ snRNA-seq data with data from two previously published MS studies^14,15^ (**Supp. Fig. 3**). Aligning the FPA-μFTIR spectral cluster maps with the Visium data confirmed concordance between the biochemomic, proteomic and transcriptomic data, demonstrating that the different cellular niches present in SPMS lesional and peri-lesional areas can be uniquely distinguished by their biomolecular profiles.

**Figure 1.**
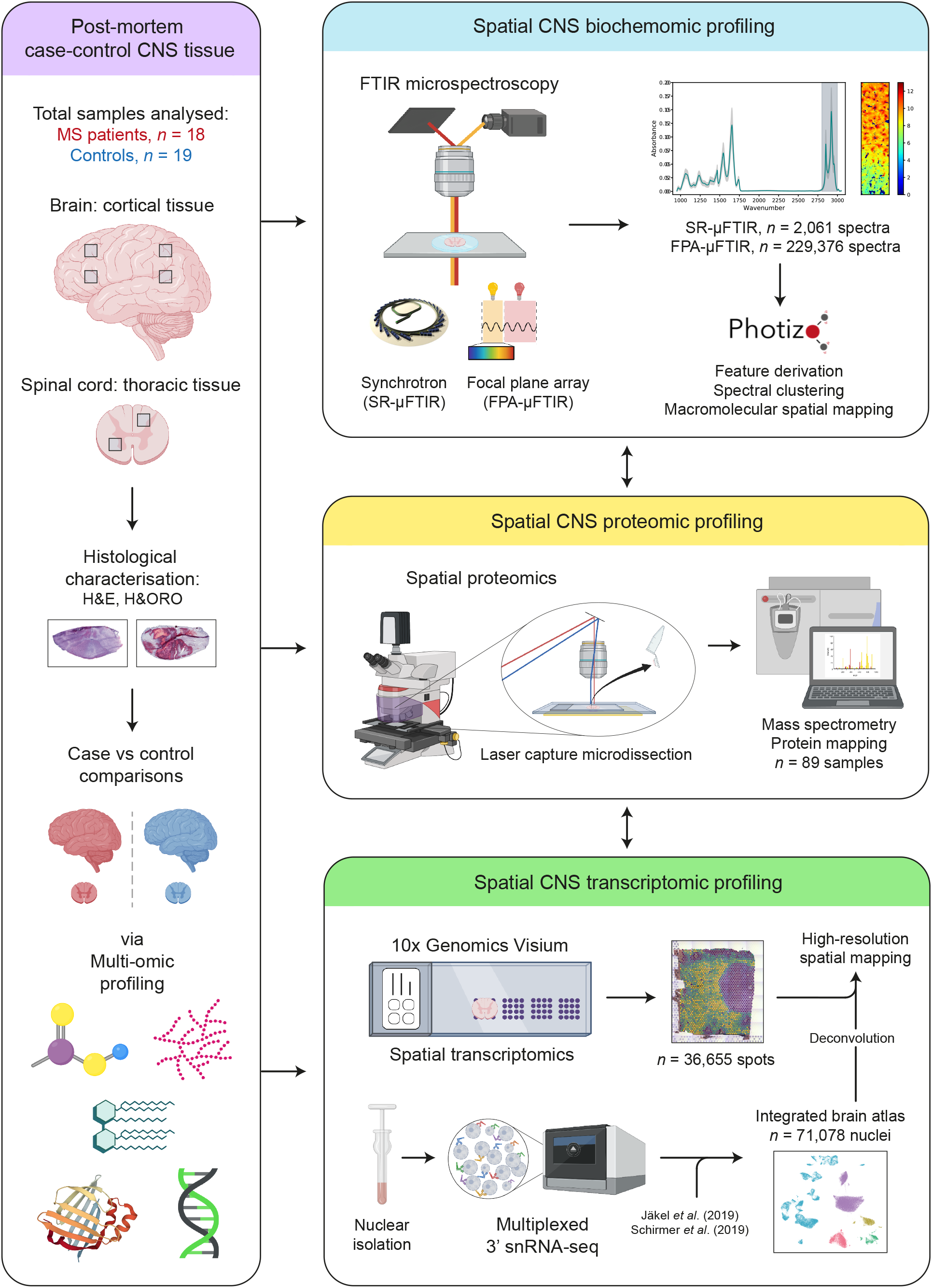
A high-resolution, spatial multi-omic atlas of progressive MS. Spatial biochemomic, proteomic and transcriptomic workflow for the analysis of a total of samples from 18 MS patients and 19 non-MS/non-neurodegenerative disease controls.

**Figure 2.**
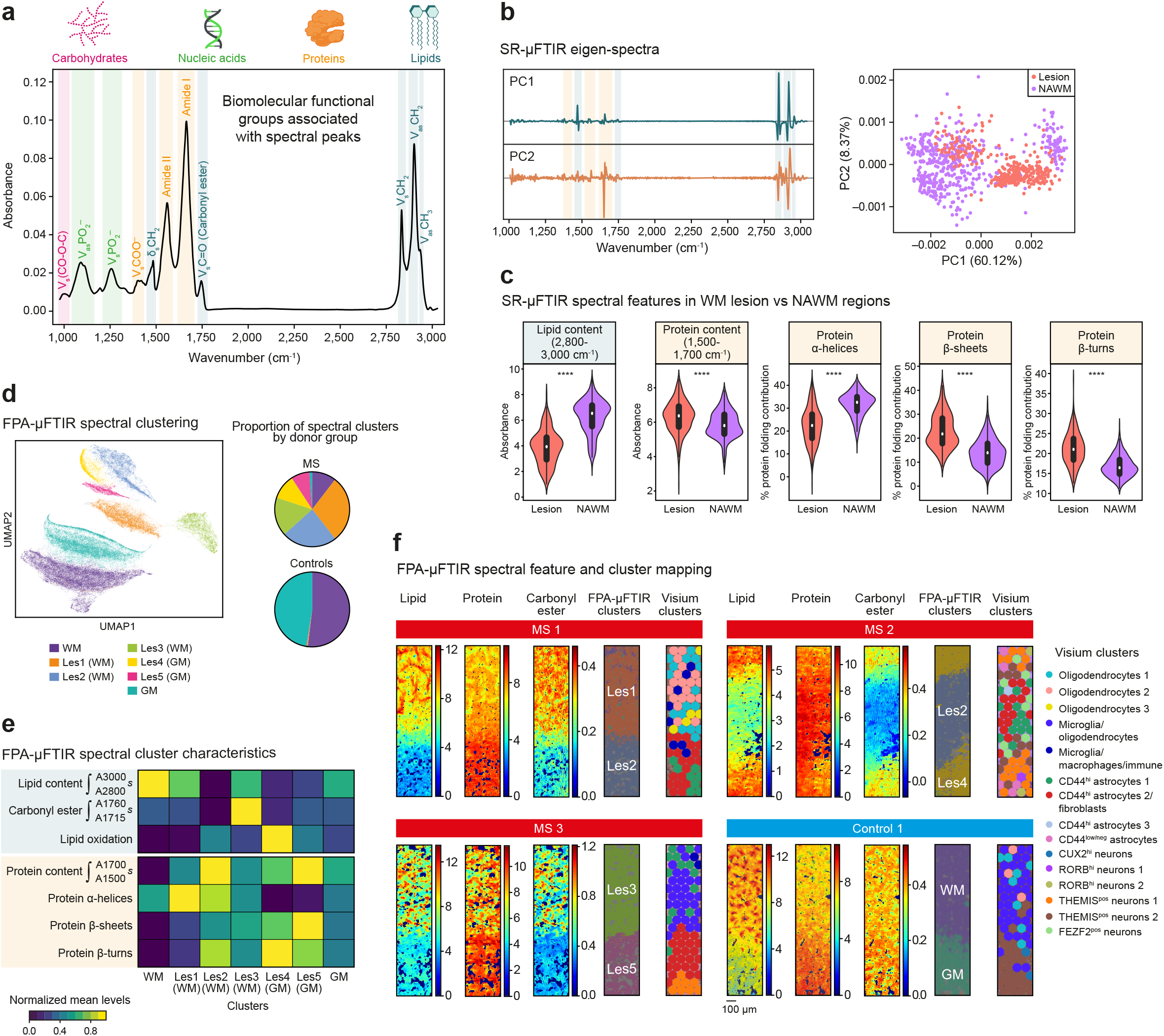
μFTIR analysis of progressive MS and control CNS tissue. **a**, Schematic representation of the functional groups and biomolecules associated with the spectral peaks detected by μFTIR spectroscopy. **b**, PCA analysis of SR-μFTIR data (2,061 spectra) collected from the analysis of SPMS lesional and normal-appearing white matter (NAWM) tissue. **c**, Comparison of the spectral features of SPMS lesional and normal-appearing WM tissue. **d**, Clustering of 229,376 FPA-μFTIR spectra from cases and controls and the cluster distribution amongst the donor groups. **e**, Heatmap of the key characteristics of the FPA-μFTIR spectral clusters. **f**, Representative images of the spatial mapping of FPA-μFTIR spectral features and spectral and spatial transcriptomic cell clusters.

**Figure 3.**
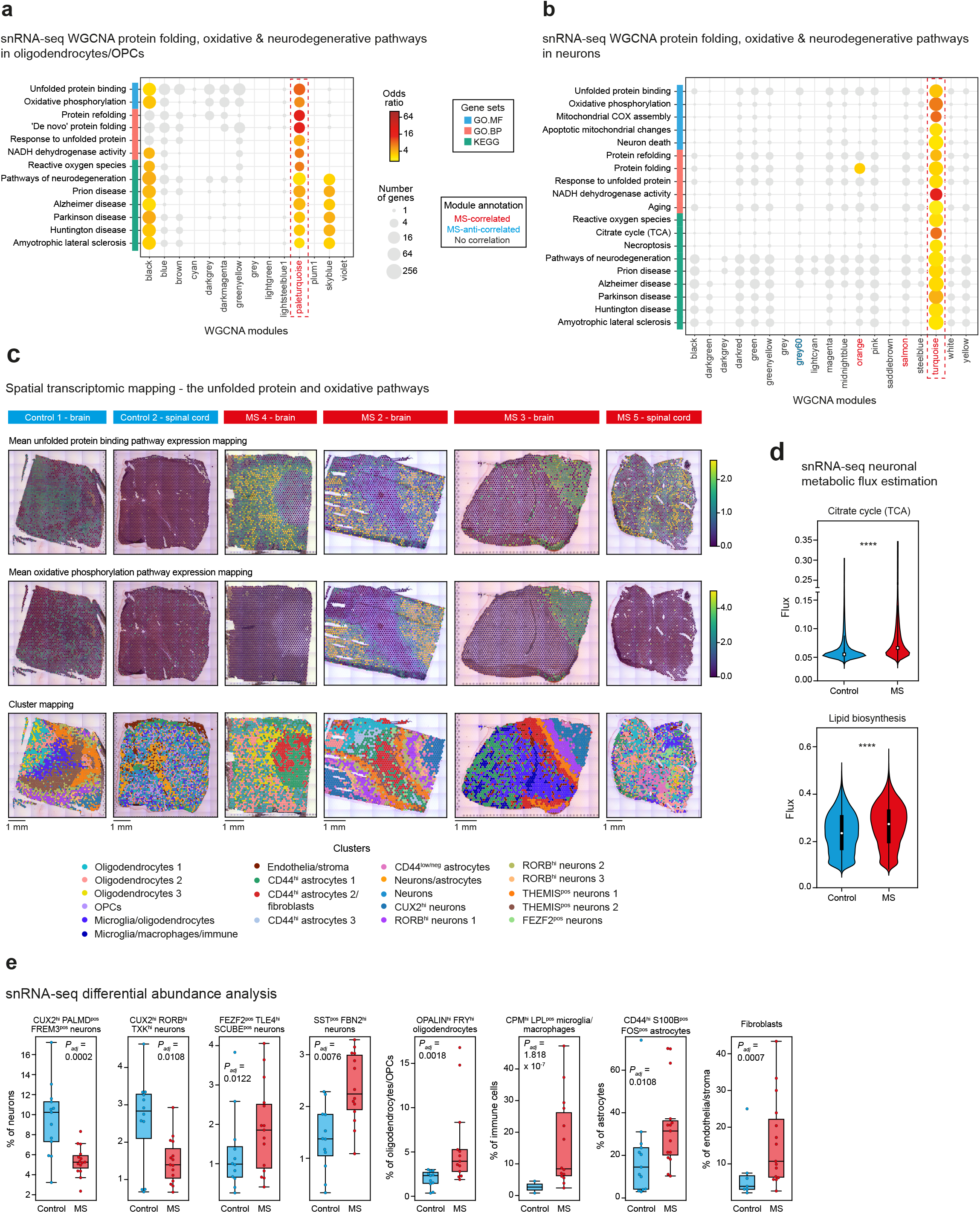
snRNA-seq and spatial transcriptomic analysis of neurodegenerative pathways. **a**, Pathway enrichment analysis of weighted gene correlation network analysis (WGCNA) modules from oligodendrocyte/oligodendrocyte progenitor cell (OPC) snRNA-seq. **b**, Pathway enrichment analysis of WGCNA modules from neuronal snRNA-seq. **c**, Representative images of spatial transcriptomic mapping of aberrant protein folding and oxidative stress pathways. **d**, Comparison of neuronal snRNA-seq metabolic fluxes in SPMS cases and controls. **e**, Differential abundance analysis of integrated snRNA-seq data.

Given that signatures of altered protein folding and oxidation contributed to the definition of the SPMS-specific spectral clusters, we next assessed whether we could find support for this at the RNA level. Performing weighted gene correlation network analysis (WGCNA) on the integrated snRNA-seq data to identify clusters of interconnected genes (modules), we found one SPMS-associated module (‘paleturquoise’) in the oligodendrocytes (**Supp. Fig. 4a**), and three in the neurons (‘orange’, ‘salmon’ and ‘turquoise’), as well as a disease anti-correlated module (‘grey60’) in the latter cellular compartment (**Supp. Fig. 4b**; all *P*_*adj*_<0.05). Pathway analysis demonstrated that of these, the oligodendrocyte ‘paleturquoise’ and the neuronal ‘turquoise’ modules were both enriched for pathways associated with aberrant protein folding and pathways indicative of other neurodegenerative diseases (**Fig. 3a,b**). Notably, cell-cell interaction analysis revealed *LRP1*- and *APP*-related ligand-receptor pairs amongst those interactions specific to SPMS, predominantly in the neuronal cell types and the oligodendrocytes (**Supp. Fig. 5**). LRP1 and APP have been linked to the clearance of protein aggregates and are implicated in Alzheimer’s disease and Parkinson’s disease.^16^ The oligodendrocyte ‘paleturquoise’ and neuronal ‘turquoise’ modules were also enriched for pathways associated with oxidative stress, but this was especially prominent for the neurons, where pathways relating to mitochondrial function, the citrate (TCA) cycle, aging and neuron death were additionally enriched (**Fig. 3a,b**). Scoring and mapping the presence of these pathways within the spatial transcriptomic data showed that whilst control brain and spinal cord tissue had no or minimal enrichment regardless of the cell types present in the sections, SPMS tissue was enriched for aberrant protein folding-related pathways in niches corresponding to oligodendrocytes (particularly adjacent to astrocyte-rich lesions) and neurons (**Fig. 3c**). In keeping with the FPA-μFTIR analysis and the WGCNA, oxidative pathways prominently localized to the SPMS GM (**Fig. 3c**). To confirm that the increased neuronal oxidation was linked to metabolic alterations in these cells, we performed a metabolic flux analysis. We found that consistent with the WGCNA pathway enrichment analysis, flux in the TCA cycle, which is a highly conserved process central to cellular energy production in the cell and which is also highly oxidative, was upregulated in SPMS neurons relative to controls (**Fig. 3d**). In addition, lipid biosynthesis, which serves as a source of acetyl-CoA that is the input for the TCA cycle, was significantly upregulated in SPMS neurons, indicating metabolic dysregulation as a driver for neuronal oxidative stress (**Fig. 3d**), and ultimately leading to a loss of CUX2^hi^ neuronal cell subsets (**Fig. 3e**), that are selectively vulnerable.^15^

**Figure 4.**
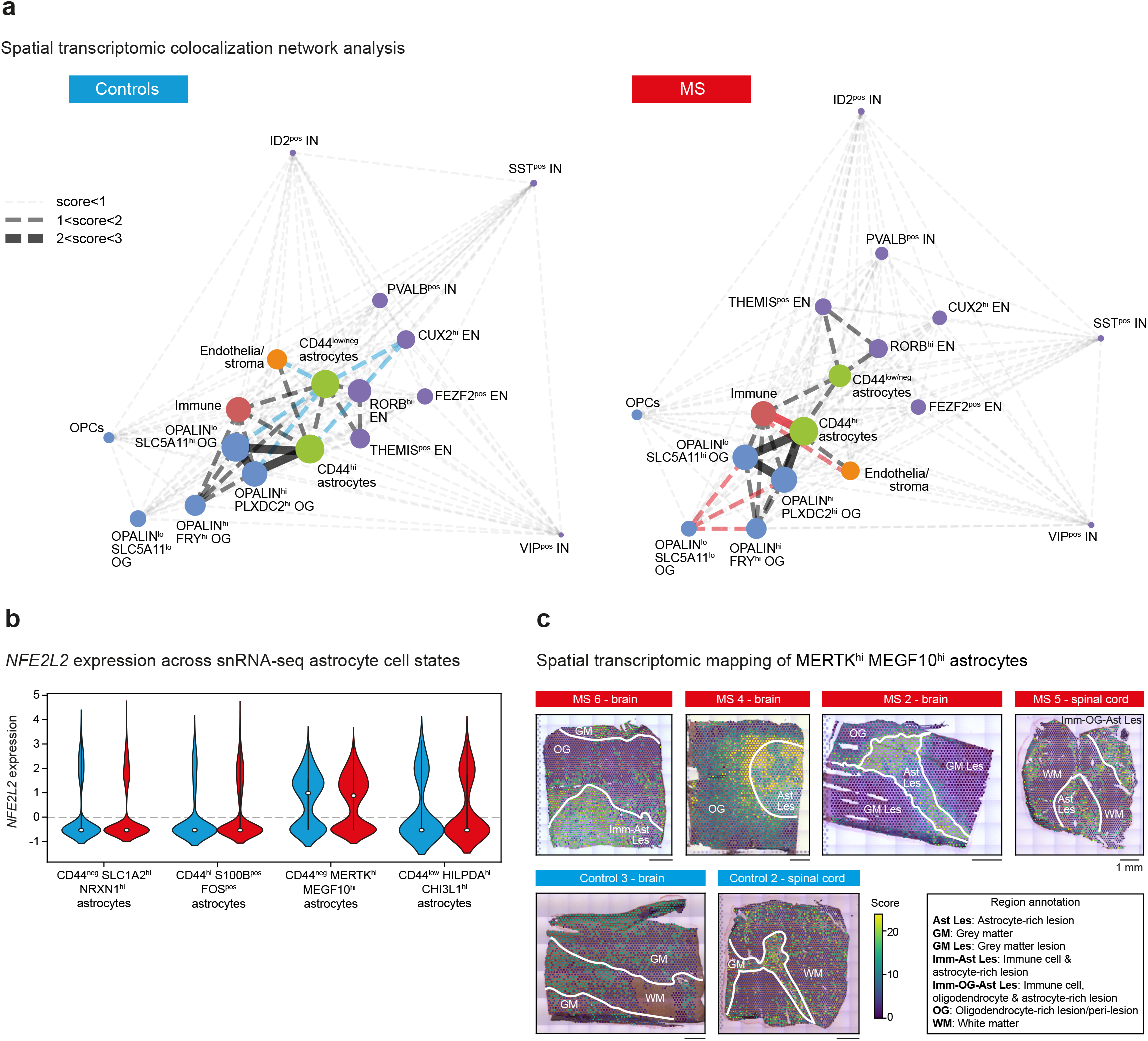
Spatial transcriptomic colocalization network analysis in SPMS cases and controls. **a**, Colocalization analysis of key cell types detected by deconvoluted spatial transcriptomics of SPMS and control CNS tissue. EN, excitatory neurons, IN, inhibitory neurons, OG, oligodendrocytes, OPC, oligodendrocyte progenitor cells. **b**, *NFE2L2* expression across astrocyte subsets in cases and controls. **c**, Representative images of the spatial transcriptomic mapping of the CD44^hi^ MERTK^hi^ MEGF10^hi^ astrocytes in case and control tissue.

Neurons, as well as oligodendrocyte progenitor cells, represent the energetic Achilles’ heel of the GM: they are simultaneously energetically demanding and have limited anti-oxidant capacity, instead relying on the metabolic support and anti-oxidant capacity of astrocytes.^17^ For example, expression of the transcription factor NRF2, encoded by *NFE2L2*, in astrocytes upregulates a neuroprotective, anti-oxidant gene program that is repressed in an astrocytic population enriched in experimental autoimmune encephalomyelitis and multiple sclerosis.^18^ Thus, we next assessed the frequency (**Fig. 3e**) and relative spatial localization of astrocyte subsets (**Fig. 4a**) in the SPMS cases and controls. We observed a significant increase (*P*_*adj*_=0.0108) in the proportion of a CD44^hi^ S100B^pos^ FOS^pos^ astrocyte population with a low expression *NFE2L2* (**Fig 3e, Fig. 4b**). This astrocyte subset was observed in WM lesions (**Fig. 3d**), and strongly colocalized with oligodendrocytes, immune cells and endothelia/stroma in SPMS; colocalization with CUX2^hi^ and RORB^hi^ neurons was reduced in cases compared to controls, which may partly be related to the disease-associated decrease in the former cell type. Intriguingly, the CD44^low/neg^ astrocytes also showed a striking change in spatial colocalization is SPMS compared to controls, despite no significant difference in their abundance. In controls these cells typically colocalize with excitatory neuronal subsets, endothelia/stroma, microglia, oligodendrocytes and other astrocytes, but in SPMS the colocalization with CUX2^hi^ neurons, endothelia/stroma and oligodendrocytes was largely lost (**Fig. 4a**). Notably, the CD44^low/neg^ astrocyte population includes the CD44^neg^ MERTK^hi^ MEGF10^hi^ cell subset which was found to have the highest *NFE2L2* expression compared to other astrocytes in both the cases and controls (**Fig. 4b**). Specifically visualizing the spatial mapping of this cell subset showed a diffuse presence of these cells in the GM and WM in controls, but a prominent localization to WM lesions and a striking absence in oxidatively stressed GM (**Fig. 4c**). These data suggest that a loss of NFE2L2^hi^ astrocytes from the GM may contribute to neuronal pathology, and that maintenance of the homeostatic astrocyte localization within the GM may be central to preventing further neurodegeneration.

In this study, we provide a high-resolution, multiomic spatial map of SPMS patient and control brain and spinal cord tissue. We observe an unparalleled capacity of the biochemome, as assayed through μFTIR, to distinguish disease-specific, pathological cellular niches. We find that these niches are underpinned by aberrant protein folding, oxidative stress and metabolic dysfunction, and a concomitant alteration in the astrocytic architecture across the WM and GM, that may be particularly critical for neurons and that may have important therapeutic implications. For example, our findings may be consistent with the observed capacity of siponimod to reduce risk of disability progression in SPMS, given its impact on blocking astrocyte activity and reducing astrogliosis.^19^ Conversely, whilst the NRF2-activating drug dimethyl fumarate is approved for relapsing-remitting MS,^20^ trialling in progressive disease has not been efficacious,^21^ perhaps due to a need for a restoration of the GM astrocytic architecture in order for the NRF2 activation to exert its protective effect on the neuronal compartment.

## Author contributions

MGP and CAD conceptualized the study and wrote the manuscript, and CAD supervised the work. MGP performed all experiments and all computational analyses. CRG provided the single-cell and spatial transcriptomics pipelines and contributed to the data analysis. HYY contributed to all experiments. TT contributed to single-cell data analysis. SD and RD contributed to proteomics experiments and analysis. MA and GC contributed to spectroscopic experiments. JF contributed to histological analysis.

## Acknowledgements

This work was performed with support from the Wellcome Trust and Royal Society (204290/Z/16/Z) to CAD, and the Interdisciplinary Bioscience DTP, supported by the BBSRC to MGP. Tissue samples and associated clinical and neuropathological data were supplied by the Multiple Sclerosis Society Tissue Bank, funded by the Multiple Sclerosis Society of Great Britain and Northern Ireland, registered charity 207495.

## Conflict of interest

The authors declare that they have no competing interests.

## Methods

### Human donor tissue

Postmortem fresh-frozen human brain and spinal cord samples were obtained from the UK Multiple Sclerosis & Parkinson’s Disease Tissue Bank at Imperial College London, under the Multicentre Research Ethics Committee approval (reference number 08/MRE09/31+5). Samples had a mean RIN value of 6.95 and were initially characterised with haematoxylin and Oil-Red-O staining. Briefly, 10 μm tissue sections were fixed in 10% formalin, slides were washed and dipped three times into 60% triethyl phosphate solution. Slides were immersed in Oil-Red-O solution for 15 min. Excess dye was washed by dipping slides three times in 60% triethyl phosphate and washed in water. Samples were then counterstained with haematoxylin for 5 min and subsequently washed in water for 5 min, air dried, mounted with 50% glycerol and imaged in an Aperio S2 slide scanner with 20X magnification.

### μFTIR sample preparation

Ten μm-thick tissue sections were placed on calcium fluoride infrared (IR) optical windows (Crystran). They were then fixed using 4% paraformaldehyde (PFA) at room temperature for 10 min and washed 3 times with water to ensure removal of any remaining PFA. Samples were then air-dried and stored at room temperature.

### SR-μFTIR

Data collection was performed at the MIRIAM B22 beamline at Diamond Light Source (Didcot, UK). Image data were collected at 36X magnification on a Hyperion 3000 microscope with a 15×15 μm^2^ aperture and 5 μm scanning area overlap with neighboring collection areas for optimal spectral quality and to maximize resolution. Data were collected with a spectral resolution of one data point per 0.964 cm^−1^. Each sample area was scanned 64 times. A clean tissue-free area was also selected and scanned 64 times as a background reference.

### FPA-μFTIR

Data collection was performed at the same location as for the SR-μFTIR. Image data were collected at 20X on a Hyperion 3000 microscope. Spectroscopy data were collected using a Bruker Vertex 80V Fourier Transform IR Interferometer with a focal-plane array (FPA) detector (64×64, 40 μm pixel size, for near- and mid-IR) with 2×2 binning, and a spectral resolution capturing one data point per 1.928 cm^−1^. Maximum absorbance was kept below 1.0. A clean tissue-free area of the slide was selected for background measurements and scanned 64 times. The total area scanned for each brain sample was 398,881.3 μm^2^, where 16,384 spectra were captured (spatial resolution = 24.34 μm^2^ per spectrum). Scanned spinal cord areas varied between 0.5-2.5 times this area, depending on features of interest in the sample.

### Pre-processing and clustering of spectral data

Following data collection, pre-processing was performed including atmospheric compensation and baseline correction (SR-μFTIR - rubberband correction; FPA-μFTIR – concave rubberband correction, 3 iterations with 32 baseline points;) on OPUS (v. 7.4). The data were then exported for analysis in our custom-built Python package Photizo.^10^ The 2,250-2,400 cm^−1^ region which is associated with atmospheric CO_2,_ was excluded from analysis. Outlier spectra with lipid peak and protein peak simultaneously below 1/4 of the maximum peak were filtered. Baseline variation is a feature associated with scattering, thus all spectra with a variation in baseline superior to 5 times the average baseline variation were also filtered. Prior to filtration, visual inspection of microscopy images confirmed these regions corresponded to areas that did not contain tissue. Following filtration, each spectrum was normalised by dividing it by the norm of that sample. Preliminary analysis of spectral quality included dimensionality reduction with principal component analysis (PCA). Eigenspectra as well as averaged spectra per sample were visualized to confirm peak position in relation to previously reported spectral features of this sample type. Integration of the area below a band of interest was used as a robust measure of the biomolecular component of interest. The protein secondary structure composition of *α*-helices, *β*-sheets, *β*-turns and random coils was quantified by μFTIR as previously described^6^:

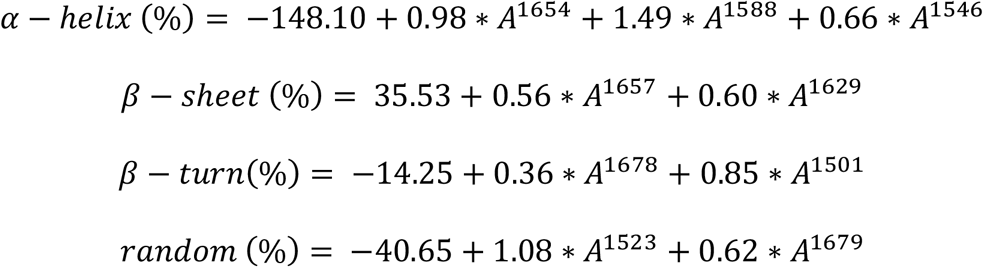

Lipid oxidation was estimated by normalising the carbonyl ester band by the total lipid content, as previously reported.^11^ For statistical comparisons of protein composition and pseudo-quantitative results from integration, a linear mixed model was used to control for scanning batch, sample effects, and spectrum inter-dependence. Features such as sex, age, cause of death and time from death to freezing did not correlate with the changes observed. All statistical comparisons were performed using Levene test for variance homogeneity, followed by the Games-Howell test.

### Spectral clustering

In order to classify tissue spectral features, we used uniform manifold approximation and projection (UMAP)^9^ to cluster the spectra. Community detection was performed with the Leiden algorithm.^22^ Spectra for each cluster were averaged for identification of cluster-distinguishing features. Cluster classification was then spatially visualized in order to correlate these results with histology.

### Spatial proteomics

Samples were sectioned for proteomic analysis onto 1.0 PEN membrane slides (Zeiss, #415190-9041-000). These were stained with H&E staining according previously reported spatial proteomics methods.^23^ In short, samples were fixed in 70% denatured alcohol, rehydrated in water and stained with haematoxylin. Tissue was subsequently washed in distilled water and stained with bluing buffer followed by another wash. Finally, the samples were stained with eosin, washed a final time and dehydrated in increasing concentrations of denatured alcohol. All samples were stored in the -80°C until next processing step.

Using a Zeiss RoboPalm LCM system, a total area of 100,000 μm^2^ was processed in each tube; microdissection was performed in 9 subareas (each area being approximately 11,111 μm^2^), since laser catapulting of tissue is more effective and precise with smaller tissue sections. For each tube, all 9 subareas were captured in 200 μl PCR tube caps with 20 μl of radioimmunoprecipitation assay (RIPA) buffer. The number and location of captured areas was determined based on μFTIR and Visium data; 89 samples were captured in total. Following collection, the samples were frozen immediately on dry ice and stored at -80°C for <3 days until proteomic preparation.

The sample processing for spatial proteomics was performed as previously reported.^23^ Briefly, the RIPA buffer containing the microdissected samples was thawed for 30 minutes followed by centrifugation. To ensure maximum protein capture, the caps were rinsed, followed by a second centrifugation. Benzonase was added and left to incubate at room temperature for 30 minutes, followed by dithithreitol (DTT) at a final concentration of 5mM was added and tubes were incubated for 30 minutes. Iodoacetamide (IAA) at a final of concentration 20 mM was added and incubated for 30 minutes. Sera-Mag-A and Sera-Mag-B Bead mix was added and mixed, followed by acetonitrile (ACN; 70% final concentration), tubes were gently tap mixed and incubated for 18 minutes at room temperature for protein to bind to beads. Samples were placed on magnets for 2 minutes and supernatant was discarded. Samples were rinsed twice with 70% ethanol for 30 seconds on magnet. Beads were rinsed with 180 μl 100% ACN for 15 seconds on the magnet. A solution of trypsin in 50 mM ammonium bicarbonate was added and beads were resuspended. Samples were digested overnight at 37°C. Following digestion, samples were bath sonicated and 180 μl of ACN was added and tap mixed. This was incubated for 18 minutes off the magnet and for 2 minutes on the magnet. All supernatant was discarded and resuspended in 5μl of 2% ACN, incubated on magnet for 5 minutes and carefully transferred to vials containing 1% formic acid. Finally, samples were stored at -20°C prior to mass spectrometry.

### Mass spectrometry data pre-processing and clustering

MaxQuant pre-processed mass spectrometry data were received and downstream analysis was performed in R 3.6.2 using Bioconductor DEP package. Initial filtration steps filtered both samples and proteins: samples with <20% of proteins present were excluded; proteins represented in <65% of samples were excluded. The 65% threshold was determined based on plotting the number of samples each protein was represented in versus. the mean intensity of that protein across samples. Following filtration, the data were log transformed and median-centered. Random forest was used for imputation. Hierarchical clustering was performed to categorise samples. In order to ensure reliability and minimize random effects from the imputation processes and hierarchical clustering, imputation and clustering were performed iteratively 100 times, with cluster classification being recorded for each sample was recorded for each iteration. Our assessment determined that cluster classification was reliable, with most samples clustering in the same group in 97-100% of iterations. Cluster number cutoffs were assessed (brain k=4, SC k=5) to partition the major detected clusters. Following this, differential expression of proteins was calculated in a pair-wise fashion for each cluster, with proteins with log fold change >1 and p<0.05 being considered statistically significant. Differentially expressed proteins (DEPs) were used for heatmap cluster visualization.

### snRNAseq nuclear isolation and nuclear hashing

Nuclei were isolated with PURE Prep Nuclei Isolation Kit (Sigma-aldrich), using the manufacturer’s recommended protocol for nuclear isolation by sucrose gradient. RNAse inhibitors (4 U/μl reaction) were added to all solutions prior to each step. Multiplexing was performed by washing and hashing nuclei from each sample with one of six unique TotalSeq™-A anti-nuclear pore complex antibodies (BioLegend). Two multiplexed sequencing batches were then sequenced by the Oxford Genomics Centre (Wellcome Centre for Human Genetics, University of Oxford).

### Pre-processing, demultiplexing and clustering of snRNA-seq data

Data were aligned with CellRanger v3.1.0 using the GRCh38 genome as reference and demultiplexed with a combination of antibody demultiplexing (DemuxEM^24^) and genetic demultiplexing based on reference SNPs (Vireo^25^ and Freemuxlet^26^), for which a reference from the 1,000 Genomes Project was prepared with CellSNP^27^ for Vireo and Dsc-pileup to prepare input for Freemuxlet. Cell demultiplexing classification required two or more of these demultiplexing methods to provide the same classification for a cell to be carried forward in analysis. Based on records of antibodies paired with each patient sample, demuxEM results were used to match genetically demultiplexed samples back to patient clinical data.

Two publicly available datasets were then integrated into our analysis.^14,15^ Sample quality control, batch correction and clustering were performed, with filtering to exclude nuclei with <200 genes, >10% mitochondrial content, <700 total UMI counts, and with a doublet score>0.25. Genes present in less than 3 nuclei were excluded. Following pre-clustering, nuclei with at least two of the following features were further excluded: low total UMI counts, lack of distinguishing markers, large contributions from mitochondrial genes, high expression levels of markers from multiple major cell type groups simultaneously and higher doublet scores than average. This resulted in a total of 71,078 nuclei. Harmony^28^, BBKNN and Scanorama^29^ were compared as integration methods using local inverse Simpson index (LISI) score and by visual inspection of UMAPs. Harmony-corrected data was used for downstream analysis. 100 PCs were used for UMAP dimensionality reduction and community detection was performed with the Leiden algorithm. Major cell groups were selected based on major cell type markers and subclustered further.

### Differential abundance analysis

Cluster counts were converted to centered log-ratios (CLRs) using the ALDEx2 R package. The median CLR for each cluster-sample combination was used for PCA which was performed using prcomp (R version 3.6.2) with default parameters. ANOVA was then used to identify confounding features contributing to top principal components; the study from which each snRNAseq data set was generated was identified as a contributing factor, and was therefore controlled for in the composition model as a covariate (‘meta_experiment’). Cell counts (prefiltered using the same criteria as for the PCA) were modelled using the glmQLFit function from edgeR with disease status and meta_experiment included as covariates. Differential abundance testing was performed using the quasi-likelihood F-test, and p-values were corrected using the Benjamini-Hochberg adjustment. Unlike our study and the Schirmer data set^15^, the Jäkel study^14^ only sampled white matter areas and therefore was not included in this analysis to avoiding tissue site-associated confounding. A false discovery rate cutoff (FDR)<0.1 was used.

### Weighted gene correlation network analysis (WGCNA)

Pseudobulked snRNA-seq data were batch corrected using ComBat^30^ and then the cornet package was used for WGCNA^31^. All presented p-values were adjusted using the Benjamini-Hochberg correction.

### Cell-cell interaction analysis

Cell-cell interaction analysis was performed using the Network Analysis Toolkit for Multicellular Interactions (NATMI)^32^ with the lrc2p database as reference, employing the default parameters, with communication networks having edges filtered for specificity >0.1. [18]

### Metabolic single cell flux estimation analysis (scFEA)

In order to estimate accumulation/depletion of metabolites and metabolic module flux we used filtered and normalized gene expression. The analysis of all single nuclei was performed jointly using single cell flux estimation analysis (scFEA)^33^. Metabolite estimation was compared for cases and controls within each major cell type cluster. Supermodules were estimated by the sum of relevant modules within a given supermodule. Statistical comparisons were performed with variance homogeneity testing (Levene’s test). Having identified that across metabolites and cells certain comparisons had homogeneous variance and others not, a non-parametric test (Games-Howell) was used across all comparisons in order to maintain a standardised approach which did not assume equal variance. P-values were adjusted using the Benjamini-Hochberg correction.

### Visium spatial transcriptomics

In order to obtain sections with a 6mm x 6mm area containable within the active area of the Visium slides, the tissue samples were scored using a custom blade made by the Old Road Campus Research Building workshop (University of Oxford). CNS samples were sectioned with 10 μm thickness. Sections were placed in the active area of the slide and stored at -80°C for <5 days until use. We performed the manufacturer’s gene expression protocol (protocol version CG000239 Rev A). Briefly, we stained samples with H&E and imaged on a Zeiss RoboPalm Laser Capture Microdissection system at 10X magnification. This was followed by tissue permeabilization (18 min, determined with the Visium Tissue Optimisation kit, protocol CG000238 Rev A). Reverse transcription was performed using a Thermo Fisher Scientific Applied Biosystems Verity thermocycler, followed by second strand synthesis and cDNA denaturation in 0.08 M solution of potassium hydroxide. The solution from each well was harvested and transferred into 5 μl Tris-HCl (1M, pH 7.0). PCR optimal cycle number was determined using a QuantStudio system, where 1 μl of each sample was amplified using KAPA SYBR FAST qPCR Master Mix with low ROX. The number of cycles at 25% of the RFU plateau for each sample was determined and ranged from 17 to 20 cycles. This cycle number was then used for PCR amplification of the remaining sample volume. Samples were then stored at - 20°C and until further processing and sequencing at the Oxford Genomics Centre.

### Visium spatial transcriptomics image tiling and pre-processing

Histological images collected on the Zeiss RoboPalm LCM system were tiled using FIJI package of ImageJ version 1.0.0-rc-69/1.52p; white balance correction was performed using a macro written by Vytas Bindokas; Oct 2006, Univ. of Chicago with modifications by Patrice Mascalchi. The images were then loaded and aligned in relation to fiducial frames for spatial gene expression mapping using Loupe Browser. Visium data were aligned using SpaceRanger and filtered based on total gene counts (brain: >500; spinal cord: >500, <15,000).

### Visium spatial transcriptomics deconvolution and co-localization networks

Data deconvolution was performed using cell2location^34^. Co-localisation networks were estimated using deconvoluted data normalised per spot, where the top 6 cells per spot were considered to co-localize in that spot. This was estimated for all spots and recorded in the form of an adjacency matrix which was then normalized and plotted using the Python library networkx. Importance of cell types in the matrix was estimated using the PageRank algorithm^35^.

**Supplementary Figure 1.**
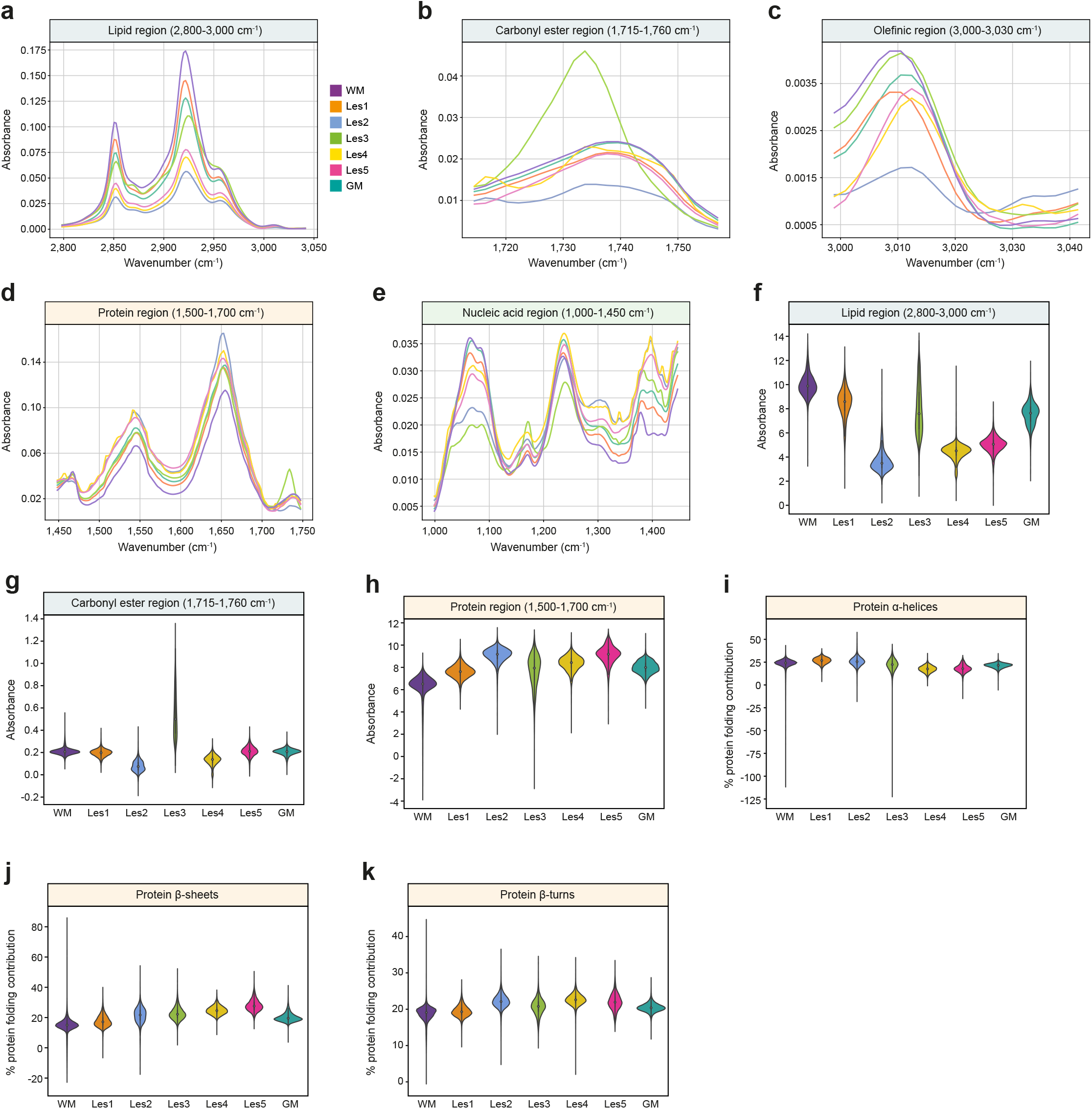
FPA-μFTIR spectral cluster features. Mean spectra of the FPA-μFTIR clusters in the lipid, carbonyl ester, olefinic, and protein regions (**a-e** respectively). Violin plots of spectra in the lipid, carbonyl ester and protein regions (f-h respectively), and of percentage of protein alpha-helices, beta-sheets, and beta-turns contributing to total protein folding (**i-k** respectively).

**Supplementary Figure 2.**
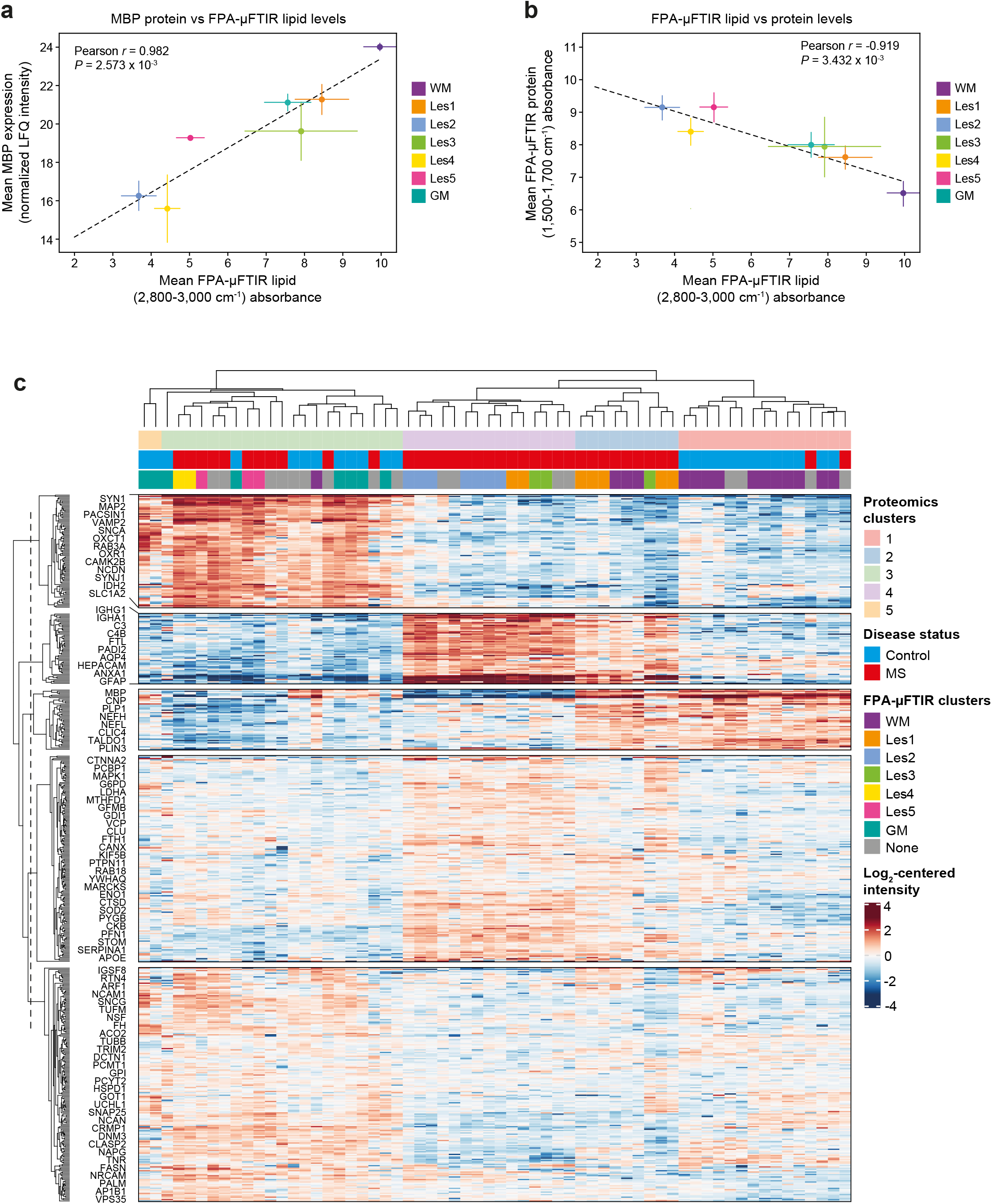
The relationship between μFTIR clusters and proteomics. **a**, Pearson correlation of the lipid absorbance (detected by FPA-μFTIR) and the myelin basic protein (MBP, detected by spatial proteomics) for the spectral clusters aligned with the spatial proteomics areas. **b**, Pearson correlation of the lipid and protein absorbances (detected by FPA-μFTIR) for the spectral clusters. **c**, Heatmap of the proteomics sample clusters and the distribution of the aligned FPA-μFTIR spectral clusters.

**Supplementary Figure 3.**
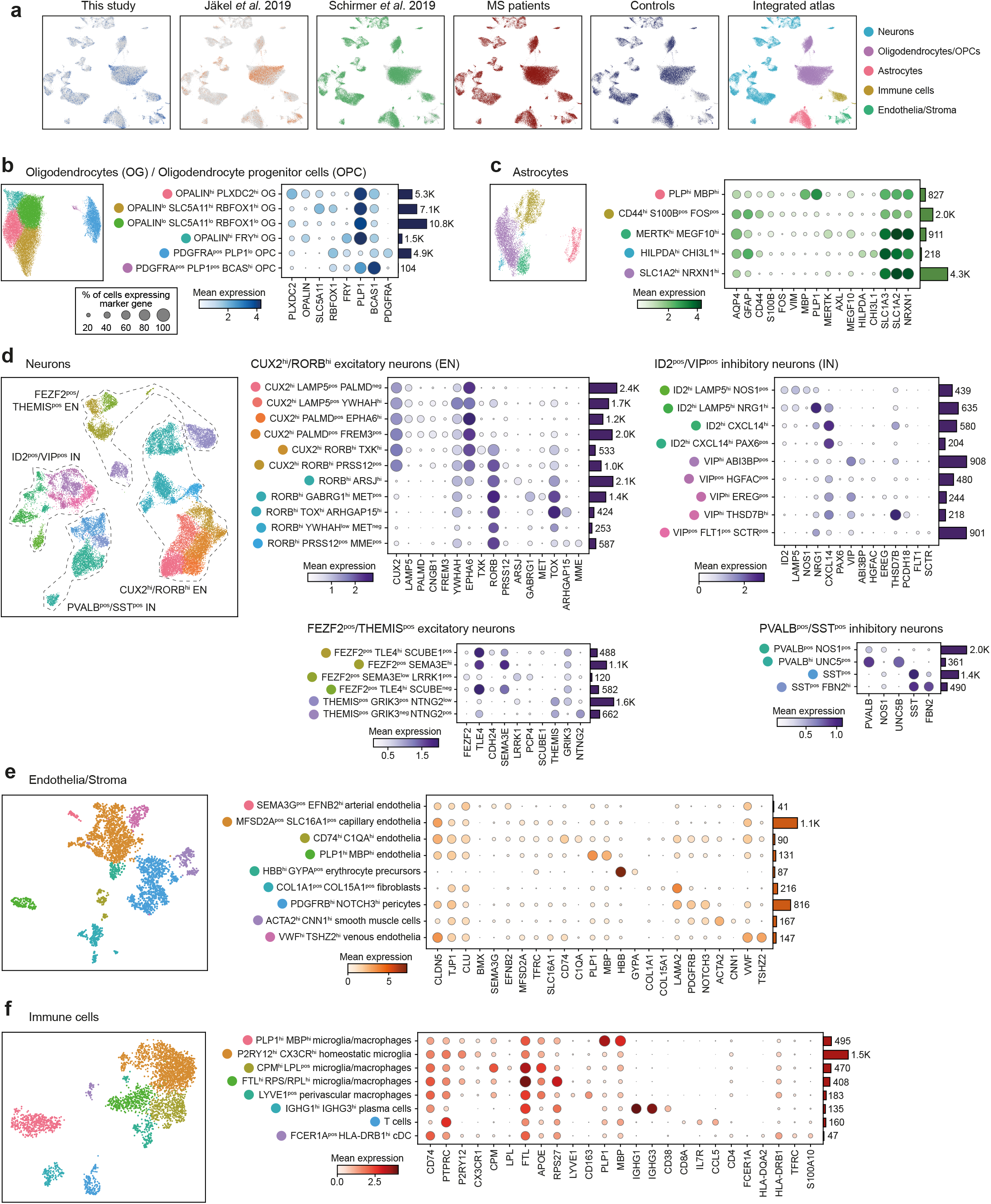
An integrated brain snRNA-seq atlas comprising 71,078 nuclei. **a**, Relative distribution of integrated nuclei by study and disease status. UMAPs and key marker dot plots for OGs/OPCs, astrocytes, neurons, endothelia/stroma and immune cells (**b-f**). The legend for the percentage of cells expressing marker genes shown in **b** applies to all dot plots in the figures. Bars to the right of dot plots indicate the number of nuclei in each cluster.

**Supplementary Figure 4.**
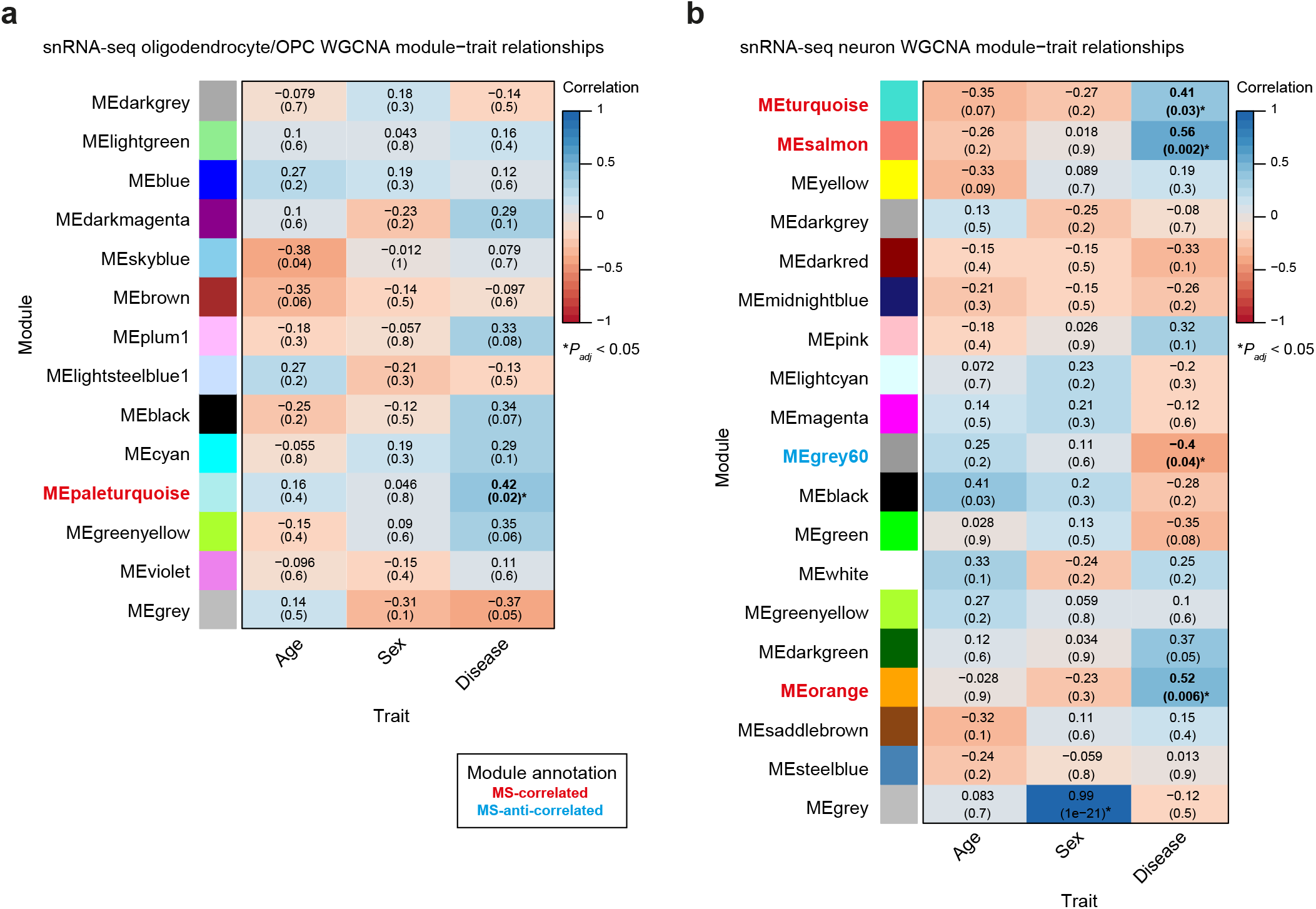
snRNA-seq weighted gene correlation network analysis (WGCNA) module-trait relationships. **a**, Oligodendrocyte/oligodendrocyte progenitor cell (OPC) WGCNA module-trait relationships. **b**, Neuronal WGCNA module-trait relationships.

**Supplementary Figure 5.**
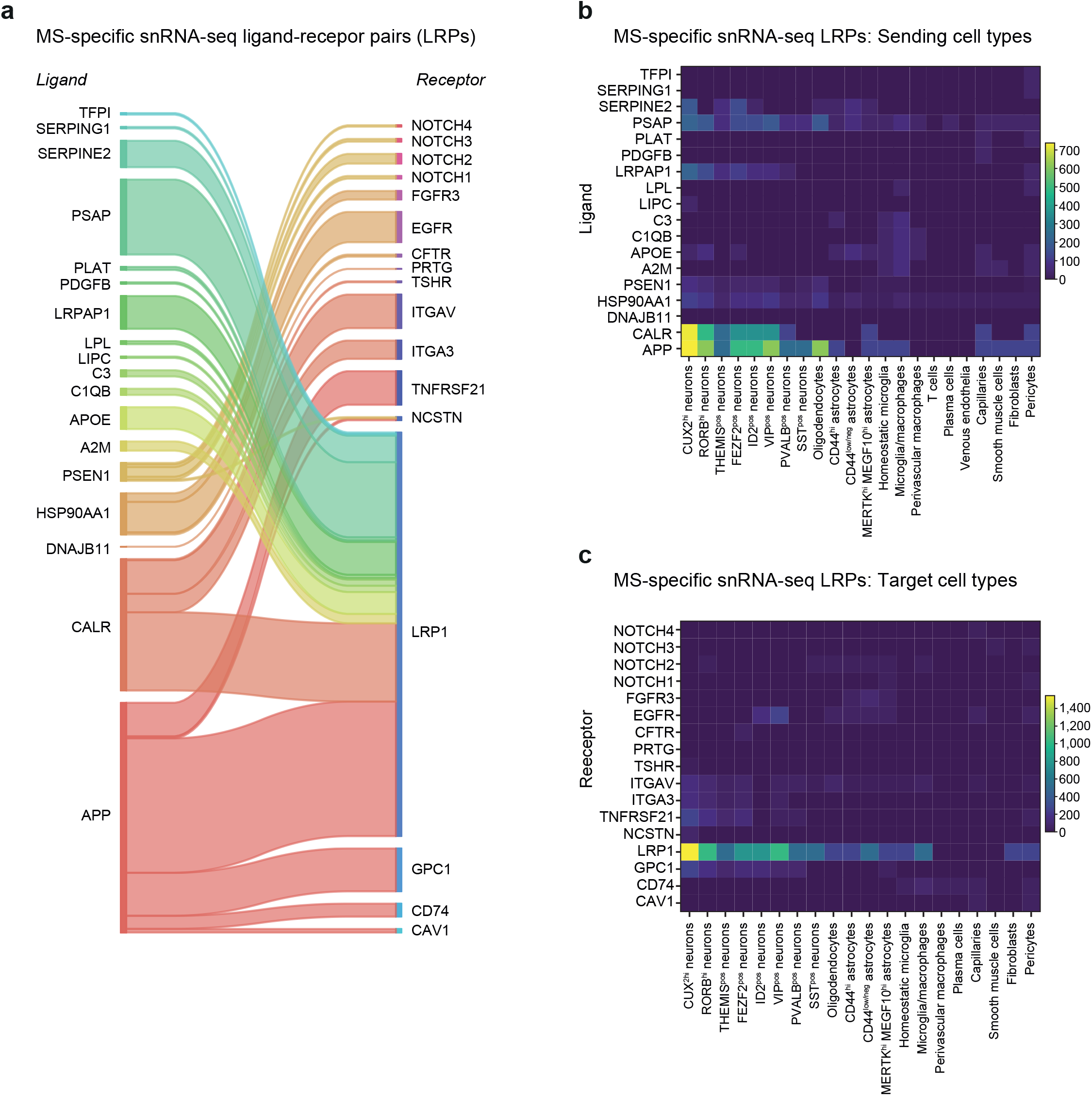
MS-specific cell-cell interactions derived from snRNA-seq data. **a**, MS-specific snRNA-seq ligand-recepor pairs. **b**, MS-specific snRNA-seq ligand expression by cell type. **c**, MS-specific snRNA-seq receptor expression by cell type.

## Notes

### Competing Interest Statement

The authors have declared no competing interest.

